# Highly Versatile, Non-Invasive Method for Collecting Buccal DNA from Free-Ranging Non-Human Primates

**DOI:** 10.1101/2020.03.29.015073

**Authors:** Aru Toyoda, Kazunari Matsudaira, Tamaki Maruhashi, Suchinda Malaivijitnond, Yoshi Kawamoto

## Abstract

Non-invasive techniques for collection of DNA samples of suitable quality and quantity are important for improving the efficiency of genetic wildlife research. The development of a non-invasive method for collection of DNA samples from wild stump-tailed macaques (*Macaca arctoides*) is described herein. Sterilized polyester rope was cut into 10 cm pieces, which were then soaked in a 20% sugar solution to bait individuals. Rope swabs were immediately collected and transferred to a lysis buffer solution after subjects had picked up, chewed, and discarded them. DNA was later extracted from the buffer. Quantitative real-time PCR and both allelic dropout and genotype failure rates were used to compare the quantity and quality of the buccal DNA samples to those of intestinal slough cell DNA samples collected from freshly dropped feces. The buccal samples yielded significantly more DNA (27.1 ± 33.8 ng/µL) than did the fecal samples (11.4 ± 15.4 ng/µL) and exhibited lower allelic dropout and genotyping failure rates for the 10 autosomal microsatellites investigated. Buccal cell collection was also simple, inexpensive, reliable, and less time-consuming compared to fecal sampling. Thus, this method should facilitate genome-wide studies of non-human primates and other wildlife species.

## Introduction

Wildlife, including non-human primates, has been subject to genetic analyses in a wide variety of research fields, such as evolutionary biology (e.g., Liu et al. 2020, Rogers et al. 2019, van der Valk et al. 2019, Williams et al. 2020), population genetics (e.g., de Manuel et al. 2016, Liu et al. 2018, Nater et al. 2017), phylogeography (e.g., Bunlungsup et al. 2016, Yao et al. 2017), pedigree analysis (e.g., Snyder-Mackler et al. 2016), and conservation biology (e.g., Lynn et al. 2016), using a variety of DNA markers. Mitochondrial DNA (mtDNA), for example, is generally used for investigating maternal relationships and phylogeography (Liedigk et al. 2015), whereas Y-chromosome genes of mammals are used to investigate paternal relationships and male dispersal (Tosi et al. 2000, Tosi et al. 2002). Meanwhile, autosomal markers, such as microsatellite and single-nucleotide polymorphism (SNP) markers, are often used to investigate population genetics and genomic diversity (Chakraborty et al. 2015, Svardal et al. 2017).

As a result of recent advances in DNA analysis technology and growing concerns over animal welfare, genetic studies of wildlife frequently use DNA samples that have been collected by non-invasive means (Lynn et al. 2016). For example, DNA samples have been collected from egg shells (herring gull, *Larus argentatus*; Egloff et al. 2009), blood-fed mosquitos (Ejiri et al. 2011), koala feces (*Phascolarctos cinereus*; Wedrowicz et al. 2013), and bug-bite blood (Sumatran rhinoceros, *Dicerorhinus sumatrensis*; Rovie-Ryan et al. 2013). DNA has similarly been collected non-invasively in genetic studies of wild, non-human primates, for example, from trapped hairs (white-headed langur, *Trachypithecus leucocephalus*; Wang et al. 2016), semen (Japanese macaques, *Macaca fuscata*; Domingo-Roura et al. 2004), urine (Japanese macaques; Hayakawa and Takenaka 1999), and saliva (mountain gorillas, *Gorilla beringei beringei*, and Grauer’s gorillas, *Gorilla beringei graueri*; Smiley et al. 2010, Chimpanzee, *Pan troglodytes*, Inoue et al. 2007). Among these DNA resources, fecal samples have been most commonly used (Chiou and Bergey 2018, Hernandez-Rodriguez et al. 2017, Orkin et al. 2020). However, fecal samples generally yield low quantities of low-quality DNA, and even though the markers used in some studies (e.g., mtDNA markers) can be amplified successfully owing to their high copy numbers (Bunlungsup et al. 2016), enormous efforts are required when examining nuclear markers (Navidi et al. 1992, Taberlet et al. 1996). One major problem with using fecal DNA samples for nuclear genotyping is allelic dropout, a phenomenon in which one of two autosomal alleles is not amplified by PCR, causing heterozygous genotypes to be misinterpreted as homozygous (Pompanon et al. 2005, Tebbutt and Ruan 2008). Allelic dropout is problematic in paternity and kinship analyses using autosomal microsatellites (Vigilant et al. 2001).

As such, development of non-invasive DNA sampling methods that allow researchers to obtain large quantities of high-quality DNA samples with low levels of contamination is needed. Buccal cell collection methods have been reported previously; such as collecting sugarcane wedges or pith of terrestrial herbaceous vegetation after their chewing by wild bonobos (Pan paniscus, Hashimoto et al. 1996, Ishizuka et al. 2018), taking oral swabs from anesthetized mountain and Grauer’s gorillas (Smiley et al. 2010), and attaching ropes to saliva-collecting devices near free-ranging Tibetan macaques (*Macaca thibetana*, Simons et al. 2012). Collecting DNA from wedges of sugar cane or other plants is a non-invasive method that does not require manipulation of animals and is thus applicable to other study sites with appropriate modification according to certain factors, such as the environment of the study and the feeding patterns of the subjects. However, methods that require specialized equipment takes time and cost to produce the device. Especially, in the wild condition, using specific devices is less flexible to collect multiple samples from several monkeys at once due to mobilities. Such methods were inapplicable to the stump-tailed macaques at our study site in Thailand because of the difficulty in preparation and storage of the bite materials. Thus, we designed alternative method for collecting buccal cells reporting here.

Herein, a non-invasive method for collecting buccal DNA samples using rope swabs is described as simple, reliable, inexpensive, and less time-consuming than other commonly used methods. To test the effectiveness of this method, two experiments were conducted. The first was a quantitative comparative test of host DNA in 41 fecal and 41 buccal DNA samples randomly selected using real-time PCR. In addition, gel electrophoresis (“gel tests”) were also used to quantitatively test DNA samples cheaply and conveniently, and their results were compared with those of costlier real-time PCR to verify their accuracy. The second experiment was a qualitative comparison based on allelic dropout and genotype failure rates in 30 fecal and 30 buccal DNA samples selected using gel tests.

## Materials and methods

### Study site

The present study was conducted at the Khao Krapuk Khao Taomor Non-Hunting Area, Phetchaburi Province, Thailand (12°47′59.2″ N, 99°44′31.1″ E), which harbors five free-ranging groups of stump-tailed macaques (*Macaca arctoides*). There are five groups: Ting group, 115 individuals; Nadam group, 91 individuals; Third group, 71 individuals; Fourth group, 75 individuals; Wngklm group, 43 individuals (Toyoda et al. 2017). The monkeys here are habituated to observer AT since 2015. This survey area is mainly a mountainous area composed of secondary forests and bamboo forests, and open areas coexist including temple and houses of local people. The moving area of monkeys was divided between north and south by large roads, and food provisioning by locals or visitors was occasionally observed along the road or at temple ground. As information on environmental conditions, mean annual temperature and annual rainfall are 27°C and 1070 mm, respectively, in the data of the nearby national park, named Keang Krachan National Park, about 30km far from this study site (Wijitkosum 2012). This site consists primarily of secondary forest, including stands of bamboo and agricultural areas.

### Collection and extraction of DNA samples

Buccal cells were collected using baited ropes (hereafter *rope swabs)*. Polyester ropes (6 mm in diameter; Takagi Corporation, Kagawa, Japan, JAN code: 4943 956 261 513) were cut into approximately 10 cm pieces, autoclaved, and dried to avoid contaminations (Figure 1). To bait individuals, the rope swabs were soaked in a 20% sugar solution (70 g cane sugar dissolved in 350 mL distilled water) for at least 30 min, and then scattered on the open ground where the monkeys were found. After being chewed (Figure 2) and discarded by a monkey, the rope swab was quickly collected and transferred to a 5 mL carrying tube containing 3 mL lysis buffer (0.5 % (w/v) in SDS, 100 mM EDTA pH 8.0, 100 mM Tris-HCl pH 8.0, and 10 mM NaCl) (Hayaishi and Kawamoto 2006). To compare the quantity and quality of the buccal DNA with that of other commonly used DNA sources, intestinal slough cells from freshly dropped fecal samples were also collected. A sterile cotton bud, which was soaked in 2 mL lysis buffer, was used to swab the surfaces of feces, following the protocol of Bunlungsup et al. (2016). To increase DNA yields, the surfaces of the feces were swabbed at least three times.

**Fig 1.**
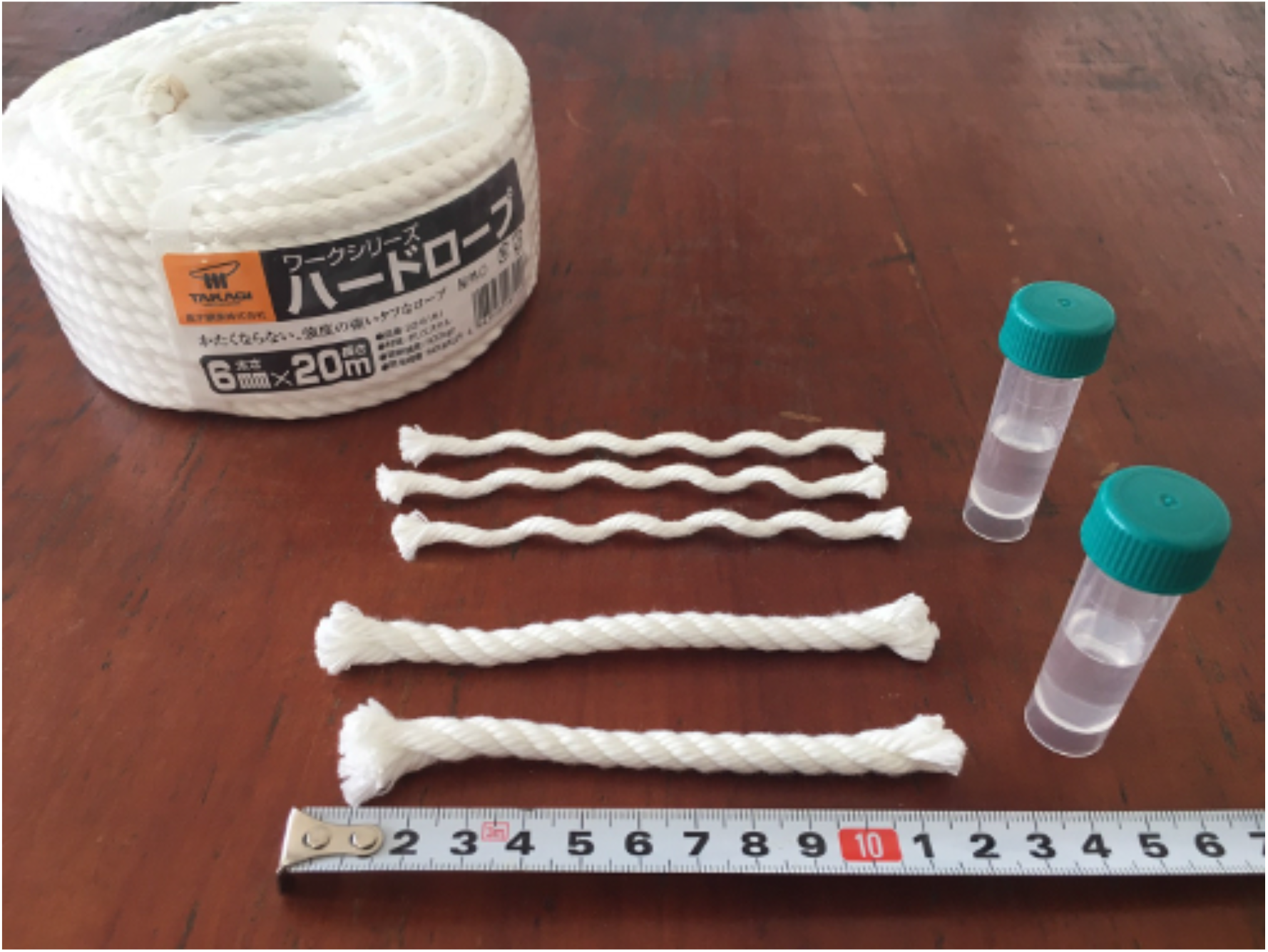
Rope swabs cut into 10cm length and 3ml of lysis buffer in 5ml tube

**Fig 2.**
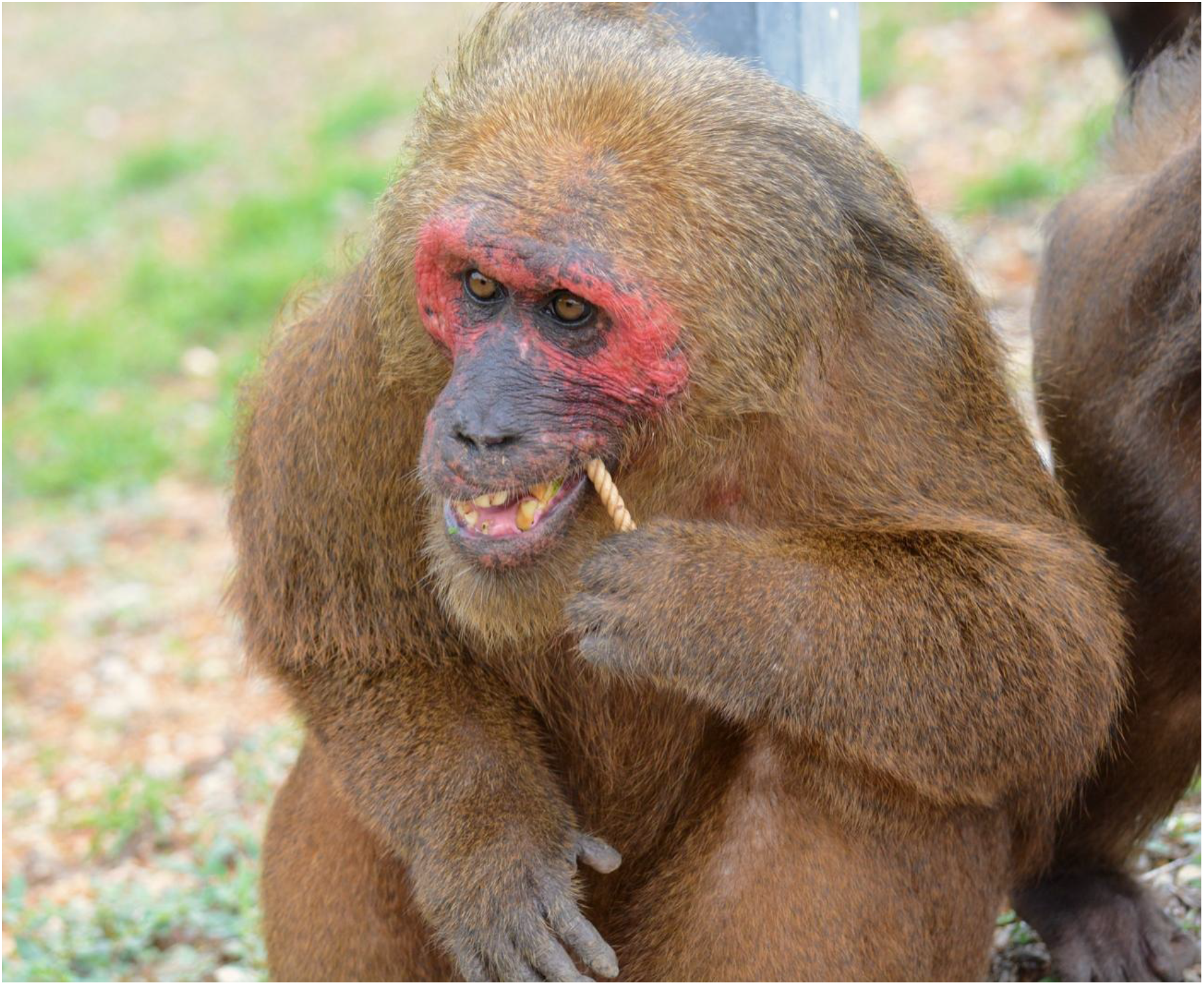
Monkey chewing a rope swab

The buccal and intestinal cells that were transferred to the lysis buffer were kept at room temperature for at least five months until DNA extraction. DNA was extracted following the procedure of Kawamoto et al. (2013). Potential PCR inhibitors were removed by adding 600 mg of hydrolyzed starch (Wako, Osaka, Japan) to 1.5 mL of lysis buffer per sample. The samples were incubated at 36 °C for 10 min, and then centrifuged at 1000 ×*g* for 15 min. Finally, 750 µL of each supernatant was processed using a commercially available DNA clean-up system (Wizard SV Gel and PCR Clean-Up System; Promega, Madison, WI, USA), and the DNA was finally eluted with 50 µL pure water. This studies including fieldwork and lab work had been conducted from September 25^th^, 2015 to June 15^th^, 2017, and 74 fecal samples and 579 buccal samples were collected.

### DNA quantification

The amount of host DNA was quantified by quantitative real-time PCR (Morin et al. 2001). Forty-one DNA samples extracted from buccal and 41 from fecal samples were selected randomly from all of the extracted DNA samples. The real-time PCR method was used because both the buccal and intestinal DNA samples were contaminated with other exotic DNA sources, such as bacteria, eukaryotic parasites, and dietary materials (e.g., plants, insects, or small animals), which could not be differentiated using conventional spectrophotometry. The sequences of the real-time PCR primers and c-*myc* probe were 5’-GCCAGAGGAGGAACGAGCT-3’ (CMYC_E3_F1U1), 5’-GGGCCTTTTCATTGTTTTCCA-3’ (CMYC_E3_R1U1), and 5’-FAM-TGCCCTGCGTGACCAGATCC-TAMRA-3’ (CMYC_E3_TMV), respectively (Morin et al. 2001). Real-time PCR was performed using a StepOnePlus real-time PCR System (Thermo Fisher Scientific, Waltham, MA, USA), and each 20 µL reaction contained 2 µL DNA template, 1× TaqMan Fast Advanced Master Mix (Thermo Fisher Scientific), 250 nM probe, and 900 nM of each primer. In addition, the PCR amplification conditions included an initial denaturation step of 95 °C for 20 s, followed by 45 cycles of 95 °C for 1 s and 60 °C for 20 s. Host DNA quantity (concentration) was determined using a standard curve made by a duplicate set of DNA with known quantity. The standard set was made from DNA extracted from the blood of a northern pig-tailed macaque (*Macaca leonina*) reared in the Primate Research Unit, Chulalongkorn University (Bangkok, Thailand). The DNA was quantified using a spectrophotometer and diluted to 10 ng/µL, 2.5 ng/µL, 625 pg/µL, 156 pg/µL, 39.1 pg/µL, and 9.8 pg/µL with deionized water. The mean DNA yields obtained from the buccal and fecal samples were compared using the Wilcoxon rank sum test in R Ver. 3.4.2 (R Core Team 2016).

Real-time PCR provides an accurate host DNA concentration for each DNA sample, and thus was appropriate for comparing the DNA yields of the buccal and fecal samples. However, real-time PCR analysis is expensive. Therefore, to select suitable samples for microsatellite genotyping, the usability of the 82 DNA samples was roughly screened using conventional PCR and agarose gel electrophoresis following the procedure of Kawamoto et al. (2013) and Ball et al. (2007) (gel electrophoresis). For the gel test, the c-*myc* gene was PCR-amplified in 12.5 µL reactions of 1 µL template DNA, 1× PCR Buffer for KOD FX, 400 µM dNTPs, 0.25 U KOD FX (Toyobo, Osaka, Japan), and 0.015 pM of both the forward and reverse real-time PCR primers, using the following conditions: initial denaturation step of 94 °C for 2 min, 45 cycles of 98 °C for 10 s, 58 °C for 30 s, and 68 °C for 30 s. The resulting amplicons were electrophoresed on 2% agarose-TAE gels, stained with SYBR Safe DNA Gel Stain (Thermo Fisher Scientific), and visualized using UV transilluminators to determine the intensity of the target band. To estimate the amount of buccal and intestinal DNA, a series of human placental DNA (Sigma-Aldrich, St. Louis, MO, USA) at concentrations of 500, 300, and 100 pg/µL were used as reference controls. When the luminous intensity of a PCR product was > 300 pg/µL of the control, the sample was considered to have sufficient yield template DNA for microsatellite genotyping and was used in the next step for microsatellite amplification. We used human placental DNA as a reference as following Kawamoto et al. (2013), that was different from the *Macaca leonina*’s DNA used in the real-time PCR. This was because of the difference of availability of the DNA standard in Japan and Thailand, and the difference of the species was considered not to affect the substantial results (Smith et al. 2002). The accuracy of the real-time PCR analysis and gel test screening were compared using the Wilcoxon rank sum test with continuity correction.

### DNA quality analysis

To determine DNA quality, the 30 paired buccal and intestinal DNA samples that passed the gel test were randomly selected for microsatellite genotyping. Ten microsatellite loci were amplified using a modified version of the two-step multiplex method (Toyoda and Malaivijitnnond 2018). During the first step of PCR, all microsatellite loci were amplified in a single 20 µL reaction that included 1 µL template DNA. During the second step, the 10 loci were divided into three subsets and were amplified in 12.5 µL multiplex PCR reactions that each included 1 µL of non-diluted amplicon from the first multiplex PCR reaction. The PCR thermocycling conditions were the same as those from the gel test, except that 35 cycles were used for the first PCR, and 45 for the second PCR. Allelic dropout rates and false allele rates were calculated using PEDANT Ver.1 (Johnson and Haydon 2007, available from http://sites.google.com/site/pcdjohnson/home/pedant). In the program, the results of two independent PCR products per sample per locus were used to estimate the allelic dropout and false allele rates. The allelic dropout and false allele rates of the buccal and fecal sample DNA were compared using the Wilcoxon signed-rank test (p < 0.05) in R. In addition, the genotype failure rate (a phenomenon in which the peak of an allele is detected at extremely low levels or is not detected) of each locus was calculated based on the duplicated PCR results, and the rates of genotype failure of the buccal and fecal DNA samples were compared using the Wilcoxon signed-rank test (p < 0.05) in R.

## Results

### DNA quantity

Analysis of the 82 DNA samples (41 buccal and 41 intestinal DNA samples) revealed that the buccal samples yielded significantly more host DNA (27.1 ± 33.8 ng/µL) than did the fecal samples (11.4 ± 15.4 ng/µL; *W* = 473, *P* < 0.001). Although 68% (28/41) of intestinal samples yielded concentrations less than 10 ng/µL, only 29% of buccal samples produced such low concentrations (12/41) (Figure 3).

**Fig 3.**
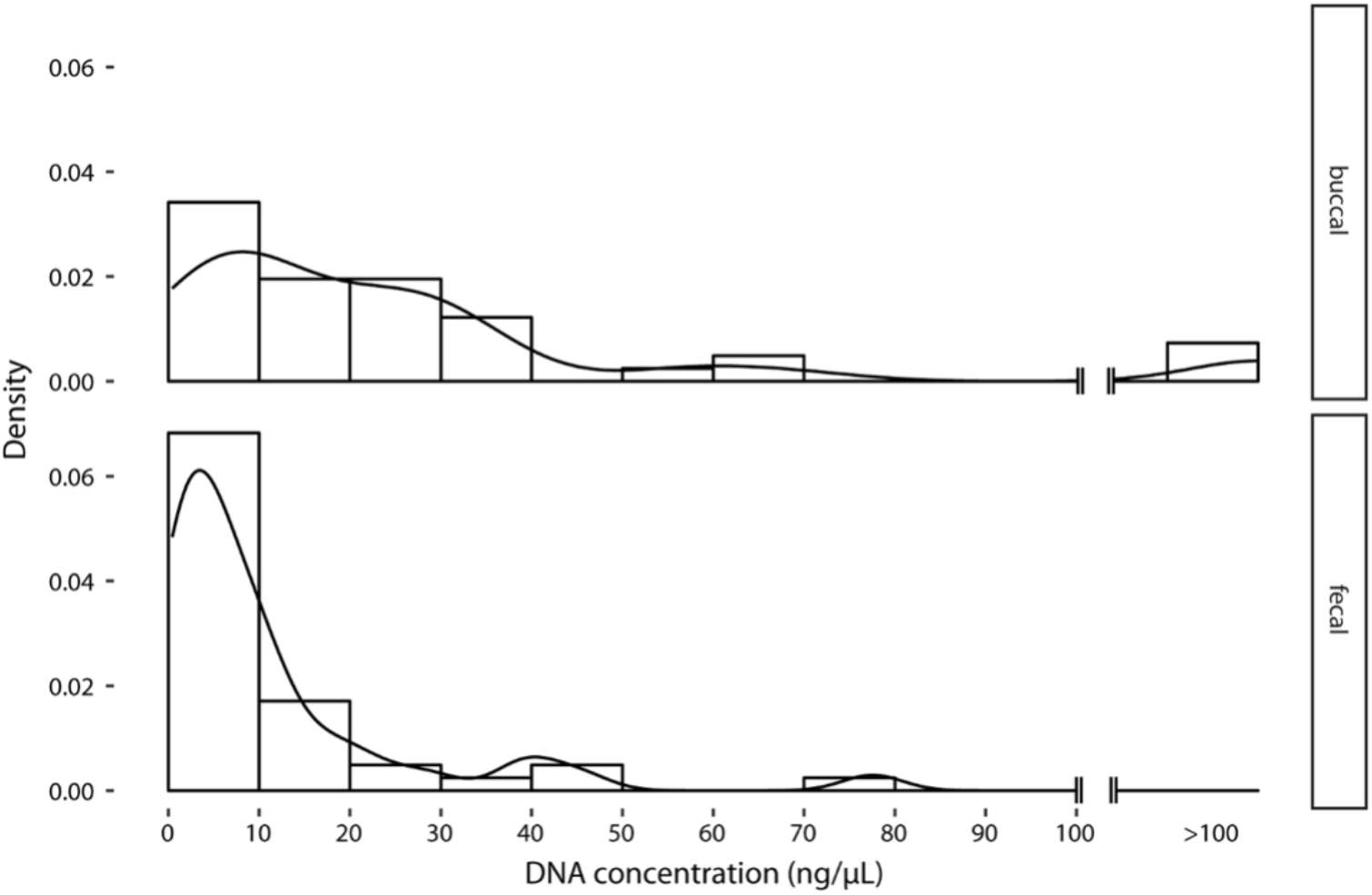
Frequency of buccal and fecal DNA in each DNA concentration zone. Although many fecal samples are dense in the low concentration zone, meaning that the sampling efficiency is not good, buccal samples shows a gentle peak overall, indicating that samples with high concentration can be more easily obtained.

The determination by the gel test was possibly made the presence/absence of the band. Of the 41 fecal and buccal DNA samples tested, 22 (53.7%) and 35 (85.4%) met the criterion for sufficient yield (≥ 300 pg/µl), respectively. The concentration of host DNA that passed and failed the gel tests as measured by real-time PCR was a significant difference (W = 991, p < 0.01), indicating that either real-time PCR or the gel test can be used for DNA quantification.

### DNA quality

For the 30 monkeys whose samples passed the gel test, the allelic dropout rate of the 10 microsatellite loci was significantly lower for the buccal (0.00%, range: 0.00 – 6 × 10^−6^ %) than for the fecal DNA samples (18.12 ± 16.12%, range: 0.00–55.96%; Wilcoxon signed-rank test, V = 44, p < 0.01; Table 1). Estimated dropout rates were used to calculate the amount of repetition necessary for accurate results at the 99.99% certainty level (Morin et al. 2001). At least 6 repetitions were needed for fecal sample analysis to produce reliable genotype data, whereas one repetition was sufficient for buccal samples.

**Table 1.** Allelic dropout and genotype failure rates of 10 microsatellite loci for fecal and buccal DNA samples of stump-tailed macaques in Khao Krapuk Khao Taomor.

| Loci | Allelic dropout rate (%) |  | Genotype failure rate (%) |  |
| --- | --- | --- | --- | --- |
|  | Fecal | Buccal | Fecal | Buccal |
| D3S1768 | 10.54 | 0.00 | 38.33 | 0.00 |
| D6S2793 | 25.00 | 0.00 | 58.33 | 2.00 |
| D7S2204 | 8.80 | 0.00 | 31.67 | 13.33 |
| D8S1106 | 13.04 | 0.00 | 45.00 | 0.00 |
| D11S2002 | 0.00 | 0.00 | 65.00 | 1.67 |
| D13S765 | 29.20 | 0.00 | 23.33 | 0.00 |
| D14S306 | 0.00 | 0.00 | 18.33 | 1.67 |
| D17S1290 | 28.45 | 0.00 | 33.33 | 5.00 |
| D18S537 | 55.97 | 0.00 | 25.00 | 3.33 |
| D19S582 | 10.21 | 0.00 | 18.33 | 0.00 |

Similarly, the genotyping failure rate had significantly lower for buccal DNA samples (2.70% ± 3.88, range: 0.0–13.3 %) than for fecal DNA samples (35.67% ± 15.35, range: 18.3–65.0%; Wilcoxon signed-rank test, V = 55, p < 0.01), although the rate was variable among the loci examined (Table 1).

## Discussion

### Advantages from sampling point of view

When fecal sample are used as genetic resources, the success in genotyping depends on the various conditions; e.g. the temperature at the time of sample collection, sample desiccation (Nsubuga et al. 2004), and salt concentration (Hofreiter et al. 2001), and skill of the collectors, as most of researcher experienced. Using the rope swab method in our study, the collection of high-quantity and quality DNA samples would be possible without much training, providing a more versatile option that is not dependent heavily on the level of experience of the sample collector. Our rope swab method may also be useful for collecting samples from infants. Indeed, our method was capable of collecting samples from infants aged 2–3 weeks, even though the feces of infants were often soft, diarrhea-like or very small and often difficult or almost impossible to collect. Thus, we strongly believe that our method would be a powerful alternative to overcome the difficulty of collecting fecal samples from infants which are indispensable for genetic analysis such as paternity test. The rope swab method is also less time consuming than fecal collection. Since the quality of DNA samples cannot be checked in-situ study, multiple fecal samples must be collected to ensure collection of an adequate sample from the target animals. On the other hand, most of buccal samples provided usable DNA, and thus, fewer specimens need to be collected from each animal. Additionally, to collect fecal samples, researchers must patiently follow the targeted animals until they defecate, which is time-consuming. Therefore, the rope swab method presented in this study has great potential to save time and mitigate these factors.

### Advantages from analysis point of view

Our study showed that the rope swab method is more effective, in terms of both quantity and quality of recovered DNA, compared to extraction from fecal samples. The rope swab method yielded up to 2.4 times more host DNA than did fecal samples and exhibited much lower allelic dropout and genotype failure rates, indicating that our method possibly facilitates genotyping analyses with fewer repetitive PCR trials, which could save time, labor, and money. This is because low DNA quantity increases genotyping errors that affect the reliability of genotyping in microsatellite analysis (Taberlet et al. 1999), and thus repeating experiments for each locus and extract is recommended (Goossens et al. 1998).

### Important notice using rope swab method

Although our method would be useful, there are several cautionary notes while collecting samples. Firstly, in the initial phase, monkeys may not chew on the rope swabs. In this case, a habituation period using fruit juice instead of sugar water to increase the attractiveness of the swab rope is recommended. From experience, however, it seems better to switch to sugar water during the sample-collection phase. Genotyping results were not stable when using DNA samples collected with orange juice, probably due to the acid or other chemical compounds present in the fruit juice.

Secondly, the collection of samples shortly after monkeys have consumed food should be avoided, especially at provisioned sites or when targeting captive animals, as fruits are the main food items given and contain acids or other chemical compounds that may inhibit PCR. Complex polysaccharides possibly originating from vegetable material in the diet are also considered potential PCR inhibitors (Monteiro et al. 1997). Thus, time of sampling may affect the quality of the sample rather than the duration for which the monkey chews the rope.

Thirdly, adjustments to the soaking time of the rope swab in the sugar-water solution and the concentration of sugar according to the condition of the subject animals or study site may be needed. Extended soaking times or high sugar concentrations could encourage monkeys to chew the rope swabs for longer periods, which may lead to greater DNA yields. However, the potential downside of a longer chewing period is that the target monkeys may move while chewing, making retrieval of the rope swabs more difficult for the researcher. Although some individuals spent significant time chewing the swabs and occasionally broke them into small fragments, no monkeys accidentally ate the rope swabs during this study period, demonstrating their safety in application.

Fourthly, the rope swabs should be well-distributed among the troop, otherwise higher-ranking males will take multiple ropes at once. When samples from subordinate individuals are needed, spreading the rope swabs over a wide area to attract high-ranking individuals, and then casting some swabs to the target individual may be an effective strategy.

Lastly, because this method requires that the rope swabs be provided to the animals, it may not be suitable for use with non-habituated, wild animals. This method also cannot be used in research sites where access to wildlife or provisioning is prohibited. Since this method involves material once contained in the mouths of animals, researchers must be aware of the possibility of touching saliva to prevent zoonosis (e.g., Kelesidis and Tsiodras 2010). When conducting behavioral observation at the same time, the possibility of influencing the behavior of the target animals must also be considered. Ultimately, the applicability of this method will depend on the specific needs and conditions of the research.

Furthermore, we must note about the standard range of quantitative real-time PCR. In this study, the standard range of quantitative real-time PCR could not cover the sample concentration range due to the fact that the quantity of DNA was extracted at a higher concentration than our assumption. We followed the protocol of Wizard SV Gel and PCR Clean-Up System and used 50 µL of water for the final elution step, though 200 µL is used in Morin et al. (2001). This difference of the final elution volume should have resulted in the higher concentration of DNA both from buccal and fecal samples in our study.

### Future possibility of application

The successful DNA collection and genotyping of *M. arctoides* using our method can be further applied to different conditions as long as researchers pay attention to risks and take precautions. For example, for populations kept in captive conditions at research institutions or individuals kept in cages in laboratories is the best condition. Also, for provisioned or well-habituated free-ranging primates such as populations living near temples which are widely seen in most Southeast Asian countries. This is a very useful method for researchers who have to obtain samples from specific individuals in a limited research period in the wild. Furthermore, with some modifications, this method can be applied for hormone and veterinary analysis (e.g., detecting a specific virus in the saliva; Musso et al. 2015, Huff et al. 2003). The non-invasive buccal cell collection method described by this study may further facilitate animal population genomic studies in both captive and field environments. Further integration of genetic information with behavioral and ecological data is expected to provide more insights into *M. arctoides*, including genetic structure and socioecological characteristics such as reproductive strategy and kinship structure.

## Acknowledgments

We are grateful to Warayut Nilpaung, Chuchat Choklap, the superintendent of the Khao Krapuk Khao Taomor Non-Hunting Area. Phanlerd Inprasoet, Wanchai Inprasoet, and Napatchaya Techaatiwatkun for providing a great deal of support in the continued success of this fieldwork; Dr. Yuzuru Hamada, Dr. Ikki Matsuda, Dr. Hiroki Koda, Dr. Ikuma Adachi, and Dr. Takeshi Nishimura for their supports in our research; the two reviewers for their valuable advice. This study was funded by the Japan Society for the Promotion of Science KAKENHI (#16J0098 to AT), a Young Science Explorer Grant from the National Geographic Foundation for Science and Exploration - Asia (#Asia-22-15 to AT), Kyoto University Foundation (to AT), and the Cooperation Research Program of the Wildlife Research Center, Kyoto University (#2015-Jiyuu-8 to AT).

## References

Ball MC, Pither R, Manseau M, Clark J, Petersen SD, Kingston S, Morrill N, and Wilson P (2007) Characterization of target nuclear DNA from faeces reduces technical issues associated with the assumptions of low-quality and quantity template. Conservation Genetics 8:577–586

Bunlungsup S, Imai H, Hamada Y, Gumert MD, San AM, Malaivijitnond S (2016) Morphological characteristics and genetic diversity of Burmese longtailed macaques (Macaca fascicularis aurea). American Journal of Primatology 74:441–455

Chakraborty D, Ramakrishnan U, Sinha A (2015) Quaternary climate change and social behavior shaped the genetic differentiation of an endangered montane primate from the southern edge of the Tibetan plateau. American Journal of Primatology 77:271–284

Chiou KL, Bergey CM (2018) Methylation-based enrichment facilitates low-cost, noninvasive genomic scale sequencing of populations from feces. Scientific Reports 8:1975

Domingo-Roura X, Marmi J, Andrés O, Yamagiwa J, Terradas J (2004) Genotyping from semen of wild Japanese macaques (Macaca fuscata). American Journal of Primatology 62:31–42

Egloff C, Labrosse A, Hebert C, Crump D (2009) A nondestructive method for obtaining maternal DNA from avian eggshells and its application to embryonic viability determination in herring gulls (Larus argentatus). Molecular Ecology Resources 9:19–27

Ejiri H, Sato Y, Kim KS, Hara T, Tsuda Y, Imura T, Murata K, Yukawa M (2011) Entomological study on transmission of avian malaria parasites in a zoological garden in Japan: bloodmeal identification and detection of avian malaria parasite DNA from blood-fed mosquitoes. Journal of Medical Entomology. 48:600–607

Goossens B, Waits LP, Taberlet P (1998) Plucked hair samples as a source of DNA: reliability of dinucleotide microsatellite genotyping. Molecular Ecology 7:1237-1241

Hashimoto C, Furuichi T, Takenaka O (1996) Matrilineal kin relationship and social behavior of wild bonobos (Pan paniscus): sequencing the D-loop region of mitochondrial DNA. Primates 37:305–318

Hayakawa S, Takenaka O (1999) Urine as another potential source for template DNA in polymerase chain reaction (PCR). American Journal of Primatology 48:299-304

Hayaishi H, Kawamoto Y (2006) Low genetic diversity and biased distribution of mitochondrial DNA haplotypes in the Japanese macaque (Macaca fuscata yakui) on Yakushima Island. Primates 47:158–164

Hernandez-Rodriguez J, Arandjelovic M, Lester J, de Filippo C, Weihmann A, Meyer M, Angedakin S, Casals F, Navarro A, Vigilant L, K€uhl HS, Langergraber K, Boesch C, Hughes D, Marques-Bonet T (2017) The impact of endogenous content, replicates and pooling on genome capture from faecal samples. Molecular Ecology Resources 2017:1–15

Hofreiter M, Serre D, Poinar HN, Kuch M, Pääbo S (2001) Ancient DNA. Nature Reviews: Genetics, 2, 353–359.

Huff JL, Eberle R, Capitanio J, Zhou SS, Barry PS (2003) Differential detection of B virus and rhesus cytomegalovirus in rhesus macaques. Journal of General Virology 84:83–92

Inoue E, Inoue-Murayama M, Takenaka O, Nishida T (2007) Wild chimpanzee infant urine and saliva sampled noninvasively usable for DNA analyses. Primates 48:156–159

Ishizuka S, Kawamoto Y, Toda K, Furuichi T (2018) Bonobos’ saliva remaining on the pith of terrestrial herbaceous vegetation can serve as non-invasive wild genetic resources. Primates 60:7–13

Johnson PCD, Haydon DT (2007) Maximum-likelihood estimation of allelic dropout and false allele error rates from microsatellite genotypes in the absence of reference data. Genetics 175:827–842

Kawamoto Y, Takemoto H, Higuchi S, Sakamaki Tart JA, Hart TB, Tokuyama N, Reinartz GE, Guislain P, Dupain J, Cobden AK, Mulavwa MN, Yangozene K, Darroze S, Devos C, Furuichi T (2013) Genetic structure of wild bonobo populations: diversity of mitochondrial DNA and geographical distribution. PLoS ONE 8:e59660

Kelesidis T, Tsiodras S (2010) Staphylococcus intermedius is not only a zoonotic pathogen, but may also cause skin abscesses in humans after exposure to saliva. International Journal of Infectious Diseases 14:838–841

Liedigk R, Kolleck J, Böker KO, Meijaard E, Md-Zain BM, Abdul-Latiff MAB, Ampeng A, Lakim M, Abdul-Patah P, J Tosi AJ, Brameier M, Zinner D, Roos C (2015) Mitogenomic phylogeny of the common long-tailed macaque (Macaca fascicularis fascicularis). BMC Genomics 16:222

Liu Z, Tan X, Orozco-terWengel P, Zhou X, Zhang L, Tian S, Yan Z, Xu H, Ren B, Zhang P, Xiang Z, Sun B, Roos C, Bruford MW, Li M (2018) Population genomics of wild Chinese rhesus macaques reveals a dynamic demographic history and local adaptation, with implications for biomedical research. GigaScience 7:1–14

Liu Z, Zhang L, Yan Z, Ren Z, Han F, Tan X, Xiang Z, Dong F, Yang Z, Liu G, Wang Z, Zhang J, Que T, Tang C, Li Y, Wang S, Wu J, Li L, Huang C, Roos C, Li M (2020) Genomic mechanisms of physiological and morphological adaptations of limestone langurs to karst habitats. Molecular Biology and Evolution 37:952–968

de Manuel M, Kuhlwilm M, Frandsen P, Sousa VC, Desai T, Prado-Martinez J, Hernandez-Rodriguez J, Dupanloup I, Lao O, Hallast P, Schmidt JM, Heredia-Genestar JM, Benazzo A, Barbujani G, Peter BM, Kuderna LFK, Casals F, Angedakin S, Arandjelovic M, Boesch C, Kühl H, Vigilant L, Langergraber K, Novembre J, Gut M, Gut I, Navarro A, Carlsen F, Andrés AM, Siegismund HR, Scally A, Excoffier L, Tyler-Smith C, Castellano S, Xue Y, Hvilsom C, Marques-Bonet T (2016) Chimpanzee genomic diversity reveals ancient admixture with bonobos. Science 354:477–481

Monteiro L, Bonnemaison D, Vekris A, Petry KG, Bonnet J, Vidal R, Cabrita J, Mégraud F (1997) Complex polysaccharides as PCR inhibitors in feces Helicobacter pylori model. Journal of Clinical Microbiology 35:995–998

Morin PA, Chambers KE, Boesch C, Vigilant L (2001) Quantitative polymerase chain reaction analysis of DNA from noninvasive samples for accurate microsatellite genotyping of wild chimpanzees (Pan troglodytes verus). Molecular Ecology 10:1835–1844

Musso D, Roche C, Nhan TX, Robin E, Teissier A, Cao-Lormeau VM (2015) Detection of Zika virus in saliva. Journal of Clinical Virology 68:53–55

Nater A, Mattle-Greminger MP, Nurcahyo A, Nowak MG, de Manuel M, Desai T, Groves C, Pybus M, Sonay TB, Roos C, Lameira AR, Wich SA, Askew J, Davila-Ross M, Fredriksson G, de Valles G, Casals F, Prado-Martinez J, Goossens B, Verschoor EJ, Warren KS, Singleton I (2017) Morphometric, behavioral, and genomic evidence for a new orangutan species. Current Biology 27:1–12

Navidi W, Arnheim N, Waterman MS (1992) A multiple-tubes approach for accurate genotyping of very small DNA samples by using PCR: statistical considerations. The American Journal of Human Genetics 50:347–359

Nsubuga, A. M., Robbins, M. M., Roeder, A. D., Morin, P. A., Boesch, C., & Vigilant, L. (2004) Factors affecting the amount of genomic DNA extracted from ape faeces and the identification of an improved sample storage method. Molecular Ecology 13: 2089–2094.

Orkin JD, Montague MJ, Tejada-Martinez D, de Manuel M, del Campo J, Hernandez SC, Di Fiore AD, Fontsere C, Hodgson JA, Janiak MC, Kuderna LFK, Lizano E, Martin MP, Niimura Y, Perry GH, Valverde CS, Tang J, Warren WC, de Magalhães JP, Kawamura S, Marquès-Bonet T, Krawetz R, Melin AD (2021) The genomics of ecological flexibility, large brains, and long lives in capuchin monkeys revealed with fecalFACS. Proceedings of the National Academy of Sciences, USA 118:e2010632118

Pompanon F, Bonin A, Bellemain E, Taberlet P (2005) Genotyping errors: causes, consequences and solutions. Nature Reviews Genetics 6:847–846

R Core Team (2016) R: a language and environment for statistical computing. R Foundation for Statistical Computing, Vienna, Austria. URL https://www.R-project.org/.

Rogers J, Raveendran M, Harris RA, Mailund T, Leppälä K, Athanasiadis G, Schierup MH, Cheng J, Munch K, Walker JA, Konkel MK, Jordan V, Steely CJ, Beckstrom TO, Bergey C, Burrell A, Schrempf D, Noll A, Kothe M, Kopp GH, Liu Y, Murali S, Billis K, Martin FJ, Muffato M, Cox L, Else J, Disotell T, Muzny DM, Phillips-Conroy J, Aken B, Eichler EE, Marques-Bonet T, Kosiol C, Batzer MA, Hahn MW, Tung J, Zinner D, Roos C, Jolly CJ, Gibbs RA, Worley KC, Baboon Genome Analysis Consortium (2019) The comparative genomics and complex population history of Papio baboons. Science Advances 5:eaau6947

Rovie-Ryan JJ, Zainuddin ZZ, Marni W, Ahmad AH, Ambu LN, Payne J (2013) Blood meal analysis of tabanid fly after it biting the rare Sumatran rhinoceros. Asian Pacific Journal of Tropical Biomedicine 3:95–99

Salgado-Lynn M, Sechi P, Chikhi L, Goossens B (2016) Primate conservation genetics at the dawn of conservation genomics. In: Wich SA, Marshall AJ (Eds) An Introduction to Primate Conservation. Oxford University Press, Oxford UK. pp. 53–77.

Simons ND, Lorenz JG, Sheeran LK, Li JH, Xia DP, Wargner RS (2012) Noninvasive saliva collection for DNA analyses from free-ranging Tibetan macaques (Macaca thibetana). American Journal of Primatology 74:1064–1070

Smiley T, Spelman L, Lukasik-Braum M, Mukherjee J, Kaufman G, Akiyoshi DE, Cranfield M (2010) Noninvasive saliva collection techniques for free-ranging mountain gorillas and captive eastern gorillas. Journal of Zoo and Wildlife Medicine 1:201–209

Smith S, Vigilant L, Morin PA (2002) The effect of sequence length and oligonucleotide mismatches on 5’ exonuclease assay efficiency. Nucleic Acids Research 30:e111

Snyder-Mackler N, Majoros WH, Yuan ML, Shaver AO, Gordon JB, Kopp GH, Schlebusch SA, Wall JD, Alberts SC, Mukherjee S, Zhou X, Tung J (2016) Efficient genome-wide sequencing and low-coverage pedigree analysis from noninvasively collected samples. Genetics 203:699–714

Svardal H, Jasinska AJ, Apetrei C, Coppola G, Huang Y, Schmitt CA, Jacquelin B, Ramensky V, Müller-Trutwin M, Antonio M, Weinstock G, Grobler JP, Dewar K, Wilson RK, Turner TR, Warren WC, Freimer NB, Nordborg M (2017) Ancient hybridization and strong adaptation to viruses across African vervet monkey populations. Nature Genetics volume 49:1705–1713

Taberlet P, Griffin S, Goossens B, Questiau S, Manceau V, Escaravage N, Waits LP, Bouvet J (1996) Reliable genotyping of samples with very low DNA quantities using PCR. Nucleic Acids Research 24:3189–3194

Taberlet P, Waits L.P., Luikart G (1999) Noninvasive genetic sampling: look before you leap. Trends in Ecology and Evolution 14:323–327.

Tebbutt S, Ruan J (2008) Combining multiple PCR primer pairs for each amplicon can improve SNP genotyping accuracy by reducing allelic drop-out. Biotechniques 45:637–646

Tosi AJ, Morales JC, Melnick DJ (2000) Comparison of Y-chromosome and mtDNA phylogenies leads to unique inferences of macaque evolutionary history. Molecular Phylogenetics and Evolution 17:133–144

Tosi AJ, Morales JC, Melnick DJ (2002) Y-chromosome and mitochondrial markers in Macaca fascicularis indicate introgression with Indochinese M. mulatta and a biogeographic barrier in the Isthmus of Kra. International Journal of Primatology 23:161–178

Toyoda A, Malaivijitnond S (2018) The First Record of Dizygotic Twins in Semi-Wild Stump-Tailed Macaques (Macaca arctoides) Tested Using Microsatellite Markers and the Occurrence of Supernumerary Nipples. Mammal Study 43:207–212.

Toyoda A, Maruhashi T, Malaivijitnond S, Koda H (2017) Speech-like orofacial oscillations in stump-tailed macaque (Macaca arctoides) facial and vocal signals. Am J Phys Anthropol 164:435–439.

van der Valk T, Gonda CM, Silegowa H, Almanza S, Sifuentes-Romero I, Hart TB, Hart JA, Detwiler KM, Guschanski K (2020) The genome of the endangered Dryas monkey provides new insights into the evolutionary history of the vervets. Molecular Biology and Evolution 37:183–194

Vigilant L, Hofreiter M, Siedel H, Boesch C (2001) Paternity and relatedness in wild chimpanzee communities. Proceedings of the National Academy of Sciences of the United States of America 98:12890–12895

Wang W, Qiao Y, Zheng Y, Yao M (2016) Isolation of microsatellite loci and reliable genotyping using noninvasive samples of a critically endangered primate, Trachypithecus leucocephalus. Integrative Zoology 11:250–262

Wedrowicz F, Karsa M, Mosse J, Hogan FE (2013) Reliable genotyping of the koala (Phascolarctos cinereus) using DNA isolated from a single faecal pellet. Molecular Ecology Resources 13:634–641

Wijitkosum S (2012) Impacts of land use changes on soil erosion in Pa Deng sub-district, adjacent area of Kaeng Krachan National Park, Thailand. Soil and Water Research 7:10–17.

Williams RC, Blanco MB, Poelstra JW, Hunnicutt KE, Comeault AA, Yoder AD (2020) Conservation genomic analysis reveals ancient introgression and declining levels of genetic diversity in Madagascar’s hibernating dwarf lemurs. Heredity 124:236–251

Yao L, Li H, Martin RD, Moreau CS, Malhi RS (2017) Tracing the phylogeographic history of Southeast Asian long-tailed macaques through mitogenomes of museum specimens. Molecular Phylogenetics and Evolution 116: 227–238

